# Imputed Genomic Data Reveals a Moderate Effect of Low Frequency Variants to the Heritability of Complex Human Traits

**DOI:** 10.1101/2019.12.18.879916

**Authors:** Kevin A Hartman, Sara R Rashkin, John S Witte, Ryan D Hernandez

**Affiliations:** Biological and Medical Informatics Graduate Program, University of California, San Francisco; Department of Epidemiology and Biostatistics, University of California, San Francisco; Department of Bioengineering and Therapeutic Sciences, University of California, San Francisco; Department of Human Genetics, McGill University, Montreal, Canada

## Abstract

The genetic architecture of complex human traits remains largely unknown. The distribution of heritability across the minor allele frequency (MAF) spectrum for a trait will be a function of the MAF of its causal variants and their effect sizes. Assumptions about these relationships underpin the tools used to estimate heritability. We examine the performance of two widely used tools, Haseman-Elston (HE) Regression and genomic-relatedness-based restricted maximum-likelihood (GREML). Our simulations show that HE is less biased than GREML under a wide variety of models and that the estimated standard error for HE tends to be substantially overestimated. We then applied HE Regression to infer the heritability of 72 quantitative biomedical traits from up to 50,000 individuals with genotype and imputation data from the UK Biobank. We found that adding each individuals’ geolocation as covariates corrected for population stratification that could not be accounted for by principal components alone (particularly for rare variants). The biomedical traits we analyzed had an average heritability of 0.27, with low frequency variants (MAF≤0.05) explaining an average of 47.7% of the total heritability (and lower frequency variants with MAF≤0.02 explaining a majority of our increased heritability over previous estimates). Variants in regions of low linkage disequilibrium (LD) accounted for 3.3-fold more heritability than the variants in regions of high LD, an effect primarily driven by low frequency variants. These findings suggest a moderate action of negative selection on the causal variants of these traits.

## Introduction

Complex traits are caused by a combination of environmental factors and genetic variants scattered throughout the genome of an organism. The mechanisms by which the alleles at those sites induce differences in traits among individuals in a population is often unknown and can be intertwined with many loci influencing many traits (Boyle, Li, & Pritchard, 2017). The collective fraction of the variance of a trait between individuals in a population that can be explained by the genetic variance between people is known as heritability, specifically the trait’s broad-sense heritability, H^2^. Family studies have measured the heritability of many complex human traits to be as high as 90% for height (Silventoinen et al., 2003), 72% for type 2 diabetes (Willemsen et al., 2015), and 83% for autism (Sandin et al., 2017).

In the search for causal loci, genome-wide association studies (GWAS) are performed (typically assuming an additive contribution of variants), and many sites have been statistically associated with a bevy of traits. However, while the collective fraction of a trait’s variance explained by the additive effects of all causal variants (the narrow-sense heritability, h^2^) can be approximated by the statistically associated variants (h^2^_GWAS_), this estimate often remains much lower than the estimates of broad sense heritability [e.g. only 16% for height (Wood et al., 2014), and 10% for type 2 diabetes (Ali, 2013)]. Even the collective fraction of variance in height explained by additive effects across all genotyped and imputed sites in these GWAS is only 60% in cohorts with n>250,000 (Wood et al., 2014). One of the potential explanations for this so-called “missing heritability” problem is the contribution of rare variants. Indeed, recent studies have implicated rare variants as a major source of missing heritability for height and BMI (Wainschtein et al., 2019)as well as gene expression (Hernandez et al., 2019), but a broader understanding of the extent to which rare variants contribute to the heritability of complex traits is needed.

The minor allele frequency (MAF) of a variant represents the frequency of the less-common allele in a sample of individuals from a population. Populations that have recently experienced rapid population growth will exhibit a larger fraction of rare alleles than populations that have not been rapidly growing. However, population genetic theory suggests that population growth alone is insufficient to drive rare variants to constitute a substantial fraction of heritability (Uricchio, Zaitlen, Ye, Witte, & Hernandez, 2016; Uricchio, 2019; Sanjak, Long, & Thornton, 2017). Natural selection is the evolutionary force that puts pressure on deleterious alleles to stay at low frequency (or be eliminated from the population) and increases the chance that advantageous alleles will increase in frequency (toward fixation in the population). If alleles that have major causal effects on a phenotype are evolutionarily deleterious, then natural selection will preferentially keep large effect alleles at low frequency, and this process can indeed drive rare variants to constitute a substantial fraction of heritability (Pritchard, 2001; Eyre-Walker, 2010; Simons, Turchin, Pritchard, & Sella, 2014; Uricchio et al., 2016). When strong effect alleles are deleterious in a population that has recently expanded (like many European and Asian populations), these evolutionary forces can act in concert to cause the genetic architecture of a trait to be dominated by rare variants (Uricchio et al., 2016; Hernandez et al., 2019; Lohmueller, 2014).

Note that a particular trait itself does not need to be under selective pressure directly to drive an effect of rare variants. If pleiotropy is common, then causal variants for a trait will have widespread phenotypic effects through interconnected networks [e.g. an omnigenic model, (Boyle et al., 2017)], and, if any one of the affected traits negatively impacts reproductive fitness, then the causal alleles could be evolutionarily deleterious. Indeed, much evidence supports the omnigenic model: 1) conserved regions of the genome tend to account for a disproportionate fraction of heritability of several complex traits (Finucane et al., 2015), 2) several attempts to infer the contribution of rare variants to heritability have found substantial evidence for it (Mancuso et al., 2016; Hernandez et al., 2019; Wainschtein et al., 2019), and 3) efforts to model the genetic architecture of complex traits as a function of purifying selection have argued that purifying selection is a prevalent force acting on causal variants (Gazal et al., 2018; Gazal et al., 2017; Zeng et al., 2018).

The primary tools for inferring narrow-sense heritability from genotypes of unrelated individuals are variance component models: Haseman-Elston (HE) regression (Haseman & Elston, 1972; Elston, Buxbaum, Jacobs, & Olson, 2000; Sham & Purcell, 2001; Bulik-Sullivan, 2015; Golan, Lander, & Rosset, 2014), Genome-based Restricted Estimation Maximum Likelihood (GREML) (Yang, Lee, Goddard, & Visscher, 2011; (Yang et al., 2010), and Linkage Disequilibrium Adjusted Kinships (LDAK) (Speed, Hemani, Johnson, & Balding, 2012). A separate category of tools, Linkage Disequilibrium (LD) Score Regression, makes use of summary statistics from genome-wide association studies to narrow-sense heritability (Bulik-Sullivan, 2015). Each approach makes assumptions regarding the genetic architecture of complex traits (such as the number of causal sites, the distribution of effect sizes, or the relationship between effect size and MAF or linkage disequilibrium), and the estimates from these techniques can be biased when models are misspecified (Evans et al., 2018; Speed, Cai, Johnson, Nejentsev, & Balding, 2017; Speed & Balding, 2019). Unfortunately, since the true underlying genetic architecture is not known in advance for a given trait, correcting for biases introduced by model misspecification may be challenging. A particularly common form of bias for variance component models is introduced when sites with different statistical properties are pooled together (e.g. heteroscedasticity). While the true causes of heteroscedasticity are often unknown, a first step to alleviate such biases is to partition sites by MAF and degree of LD (Yang et al., 2015; Evans et al., 2018). Additionally, we have noted that partitioning sites based on the MAF inferred from a larger external cohort can further reduce bias for rare variants (Hernandez et al., 2019).

The ability to study the effect of rare alleles is fundamentally limited by the difficulty and expense of accurately identifying and collecting rare variants in sufficiently large cohorts. Investigators have leveraged information from large whole genome sequencing databases such as the Haplotype Reference Consortium (HRC) (McCarthy et al., 2016) to impute millions of rare variants in cohorts of hundreds of thousands of samples [e.g. the UK Biobank (Howie, Donnelly, & Marchini, 2009; Bycroft et al., 2018)]. The UK Biobank in particular has measured a wide variety of phenotypes that we can use to ask about heritability and the genetic architecture of complex traits. However, before estimating the contribution of rare and common variants to the heritability of complex traits, we must first understand the accuracy of various inference procedures. We conducted thousands of simulations of phenotypes from genetic data and assessed how well two methods for heritability inference perform. We then explored the impact of sample size on the bias and standard errors of the estimated heritability. Lastly, we explored the impact of excluding rare MAF partitions on the inference of heritability for common variant partitions. We then applied our framework for studying variants across the MAF spectrum to infer the heritability of 72 quantitative human traits from the UK Biobank.

## Methods and Materials

### Genomic and Phenotypic Data

The primary genomic data for both simulations and the heritability inference came from the UK Biobank (Bycroft et al., 2018). The UK Biobank consists of a cohort of roughly 500,000 individuals recruited from the United Kingdom (UK) National Health Services. Individuals were recruited on the basis of age between 40 and 69 at the time of assessment. The total dataset collected included blood samples, urine samples, body measurements, self-reported ancestry, medical history, and lifestyle exposures (Bycroft et al., 2018).

The blood samples allowed the extraction and genotyping of DNA on one of two genotyping arrays designed for the UK Biobank. These genotype data were quality controlled then imputed to the HRC (McCarthy et al., 2016) with additional sites imputed to a whole genome sequence reference panel consisting of UK10K haplotype reference pane l and the 1000 Genomes Phase 3 reference panel (Chou et al., 2016). These imputed allelic dosages were retrieved as BGEN filetype (Band & Marchini, 2018). We used PLINK 2.0 (Chang et al., 2015) for further quality control and to export variants to PLINK 1 format for downstream analysis. Post-imputation quality control consisted of restricting to sites with imputation info score greater than 0.3 (Howie et al., 2009), with greater than 95% genotype hard-calls from dosage, and no deviation from Hardy-Weinberg Equilibrium (p-values > 1⨉10^-5^) (Winkler et al., 2014).

From all the individuals of the UK Biobank we applied the filtration steps described in Table 1. These filters retained a total of 366,647 high-quality, unrelated individuals. For computational reasons, we selected a subset of 50,000 of these individuals at random for both our inference of heritability and for our simulation studies. To evaluate the role of sample size, we also selected random subsets of 500 and 5,000 individuals to be used for some of the simulations.

**Table 1:** Quality Control of Genomic Data

| Quality control step | Remaining Individuals |
| --- | --- |
| Initial | 473,850 |
| Restrict to samples where self-reported and genetic sex match | 473,482 |
| Restrict to self-reported ethnicity | 445,826 |
| Restrict to samples with principal components 1 and 2 within 5 standard deviations from the mean | 440,222 |
| Remove samples with inappropriate sex-specific cancers | 440,148 |
| Restrict to samples in imputation sample file | 439,317 |
| Restrict sample those with Dish Quality Control score (DQC) $\geq 0.82$ | 439,317 |
| Restrict samples to those with hard call rates $\geq 97\%$, | 438,287 |
| Restrict to samples with heterozygosity within 5 standard deviations of the mean | 437,331 |
| Exclude at least one of any pair of individuals with 3rd degree or closer relationship (kinship $\geq (1/2)^{9/2}$), prioritizing exclusion of those with more relationships | 366,647 |

We examined all 72 quantitative phenotypes that had at least 25,000 reported values within our 50,000 person cohort. This included 42 blood measurements, 22 anthropometric traits, 5 respiratory traits, and 3 urinary traits.

### Computing MAF and LD Score Partitions

Variable sites in the 50,000 individual cohort were partitioned in two ways. First, using MAF computed using PLINK 2.0 across the full set of >360,000 unrelated, quality-controlled individuals, sites were divided into 17 MAF bins according to the following upper (closed) breakpoints: 2×10-6, 5×10-6, 1×10-5, 2×10-5, 5×10-5, 1×10-4, 2×10-4, 5×10-4, 1×10-3, 2×10-3, 5×10-3, 0.01, 0.02, 0.05, 0.1, 0.2, 0.5. We next used GCTA (Yang et al., 2011) to compute LD scores across the full set of quality-controlled individuals within each MAF bin, in sliding windows of 10 megabases along each chromosome. We then partitioned each MAF bin into high and low LD score bins using the median value LD score for that partition. This procedure resulted in a total of 34 bins of sites.

### Simulation Framework

Simulations were performed to compare two inference methods, Haseman-Elston (HE) Regression and genomic-relatedness-based restricted maximum-likelihood (GREML), as well as to identify the most suitable conditions for inference. We used PLINK 1.9 (Purcell et al., 2007) to recode the genotypes for the selected individuals into a genotype matrix, X, where the genotype of individual *i* at variant *j*(*x_ij_*) is 0, 1, or 2 copies of the non-reference allele. In each simulation, we selected a specified number of causal variants. For each causal variant, we drew effect sizes, *β_j_*, from a standard normal distribution, *β_j_* ∼*N*(0,1), with the effect sizes of the non-causal variants implicitly 0. The unscaled genetic component of the phenotype for individual *i*, *g_i_*, was then the sum of the product of the effect sizes with their corresponding genotypes 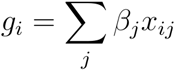,or 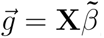. This unscaled genetic component was rescaled to give the appropriate variance, 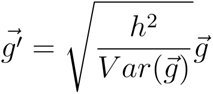, where *h*^2^ is the simulated heritability. The phenotype of individual *i*, *p_i_*, was the sum of the scaled genetic component and a residual of appropriate variance 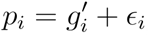, where *ɛ_i_* ∼ *N*(0,1 – *h*^2^).

In simulations where total heritability, *h*^2^, was partitioned across *m* collections of variants (or bins) as 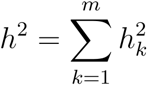, we represented each collection of variants as genotype matrices: X_1_, X_2_, …, X_m_. Letting, 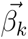, be the vector of effect sizes in collection *k* with the specified number of causal sites drawn from that partition as *β_k_* ∼*N*(0,1) and the remaining non-causal sites with effect size 0, the unscaled genetic component of the phenotype from collection *k* was 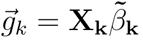, which we rescale by the appropriate heritability, 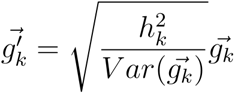. The phenotypes were then the sum of the genetic components and a residual term: 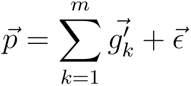, where *ϵ_i_* ∼ *N*(0,1 – *h*^2^) as before.

### Simulation Parameters

We conducted a series of 4 sets of simulations, the parameters of which are summarized in Table 2.

**Table 2:**
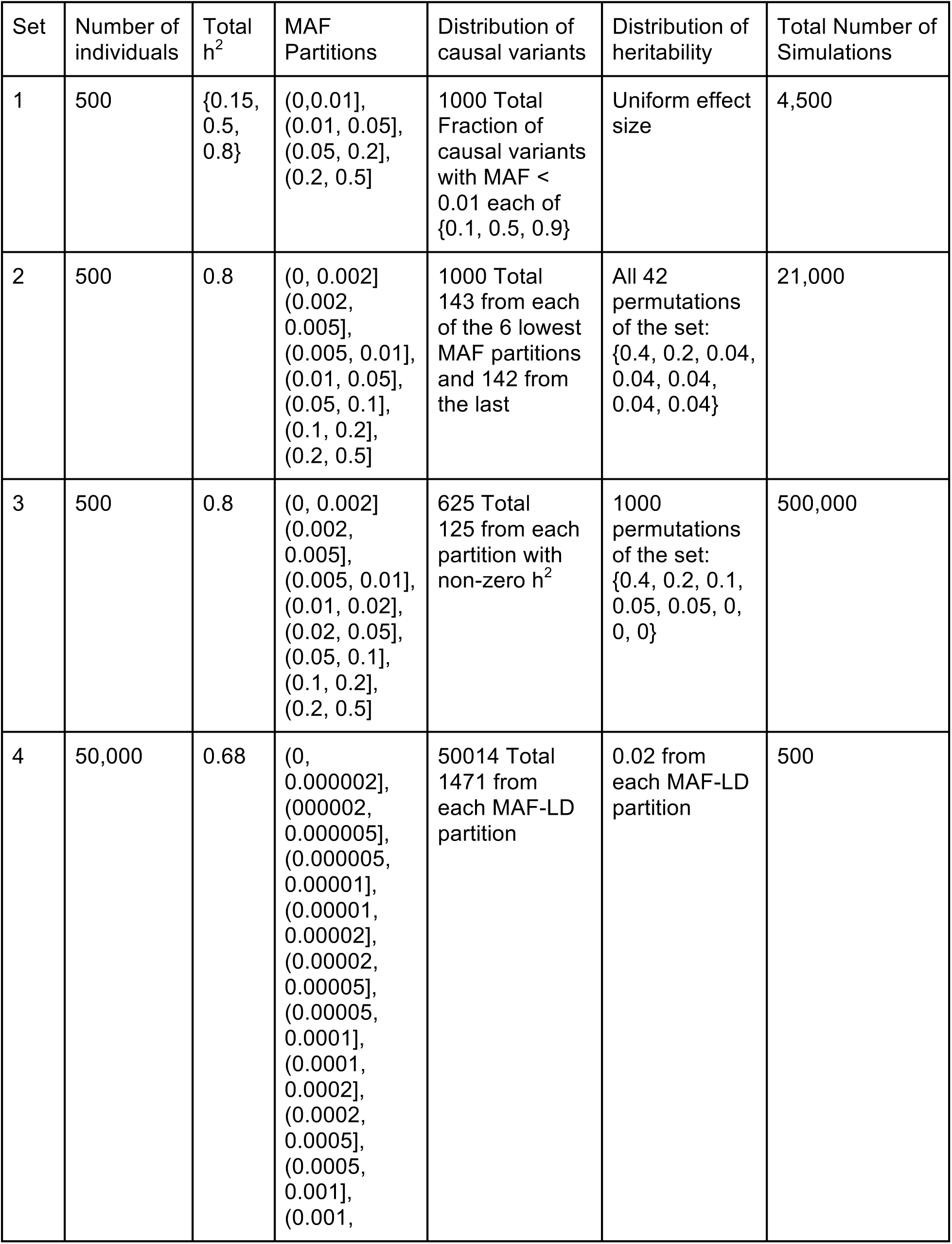
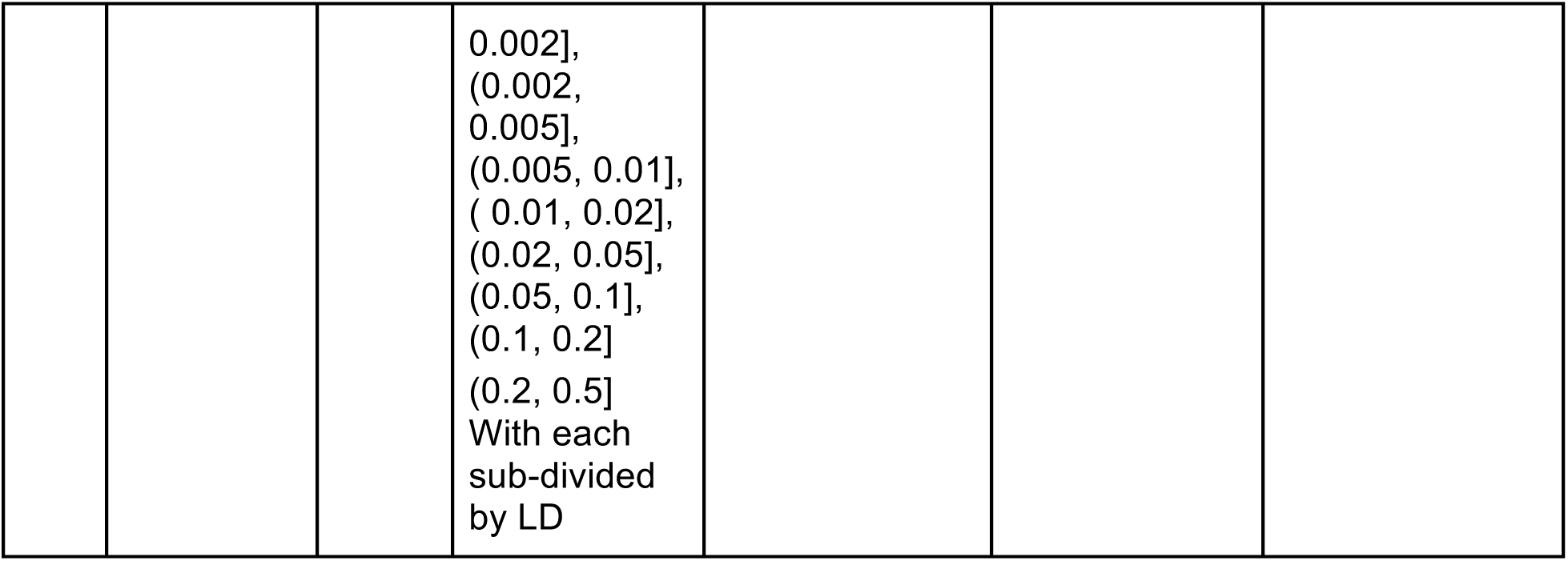
Simulation Parameters

For Sets 1-3, causal variants were drawn from the entire genome of 500 unrelated individuals. The variants were partitioned by MAF computed within the 500 individual cohort itself. In Set 1 we varied the total heritability as well as the fraction of causal variants drawn from MAF < 0.01 with uniform effect size across the MAF partitions, with 9 combinations with 500 simulations of each combination. In sets 2 and 3 we simulated 500 individuals with heritability distributed across 7 and 8 MAF partitions respectively with 500 simulations of each heritability distribution.

For Set 4, we simulated phenotypes for 50,000 individuals using genotypes from chromosomes 18-22. We partitioned these variants by their MAF in the >360,000 unrelated individuals into 17 MAF partitions. We further subdivided each of these by LD, and simulated heritability on each of the 34 MAF-LD bins, with 500 simulations.

### Heritability Inference

We used GCTA to compute the genetic relatedness matrices of individuals from the variants of each partition described. We inferred GREML heritability using GCTA’s unconstrained restricted maximum likelihood method (“--reml-no-constrain” flag) using multiple genetic relatedness matrices (GRMs). For HE heritability, we used either HE Regression as implemented in GCTA or in our own implementation in R, which we verified gave the same results to single floating-point precision. When inferring the heritability of the UK Biobank quantitative traits, we progressively included the first 15 principal components (PCs) of genetic variation and three geographic parameters of the subjects location (North-South coordinate, East-West coordinate, and distance to coast) as covariates. As the HE method of GCTA did not allow the inclusion of covariates directly, these were included as pseudo-GRMs [as per (Hernandez et al., 2019); See Supplemental Methods].

### Summarizing Inference Performance

Three metrics were used to summarize the quality of the inference for each set of simulations: bias, mean squared error, and empirical standard errors. The bias reported represents the mean of the difference between the estimated and true value.. Empirical standard errors were calculated as the average of the standard deviation of the inferred heritability for each set of simulations weighted by the number of simulations in each set.

### Software

We used PLINK1.9 v1.90b6.9 and PLINK 2.0 v2.00a2LM to manipulate the genomic data including computing MAF, filtering sites, and exporting to formats. We used GCTA version 1.92.0 to compute GRMs and to perform REML and HE regression. We used R version 3.5.1 with packages ggplot2_3.0.0, dplyr_0.8.0.1 to analyze results and generate figures. We used Python version 2.7.5 to compute covariate GRMs.

## Results

### The Role of Genetic Architecture on the Distribution of Heritability

The first set of simulations we conducted evaluated the impact of varying the fraction of causal variants that were rare (MAF < 0.01), when all variants had the same distribution of effect sizes. The distributions of these simulated heritabilities are shown in Figure 1. In this size cohort, rare variants accounted for roughly 10% of variants. Under a “neutral model” where causal variants are randomly selected from the set of all variants, ∼10% of causal variants are rare, yet they accounted for less than 1% of the simulated heritability. When we push the simulation to have an extreme excess of rare causal variants (e.g. when 90% of causal variants were rare but effect sizes maintain the same distribution across frequencies), rare variants still account for only 13% of the total heritability. These trends held regardless of total heritability (Figure S1). In all cases, the majority of heritability came from the (0.2, 0.5] MAF partition, ranging from 67% of heritability when 10% causal variants were rare to 58% when 90% causal variants were rare.

**Figure 1.**
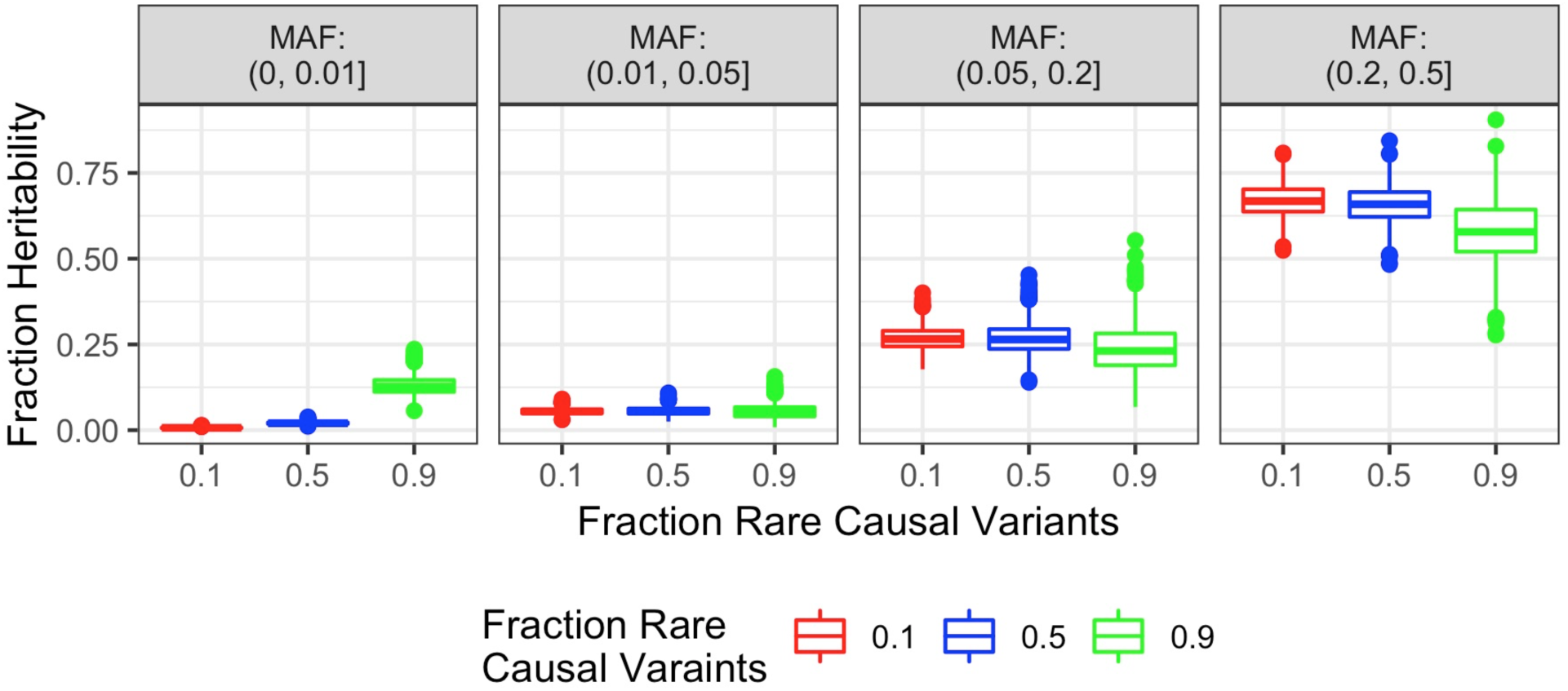
Distribution of Simulated Heritability Varying Fraction of Rare Causal Alleles. The fraction of the simulated heritability coming from different MAF partitions (horizontal panels) when varying the fraction of causal rare (MAF < 0.01) shows that under “neutral” models where variants have uniform effect sizes across the MAF spectrum, the rare variants account for very little heritability. Even when 90% of causal variants are rare, more common variants account for the majority of heritability.

Rare variants can account for a greater fraction of heritability if the distribution of effect sizes is allowed to be a function of MAF. However, the actual model relating number of causal alleles, effect size, and MAF for actual complex traits is unknown. Instead of specifying such a model and, in order to test the tools of heritability inference on the full range of possible heritability distributions, we simulated phenotypes where we directly specified the heritability coming from each MAF bin.

### Comparing HE Regression and GREML

We compared the accuracy of two common methods for heritability inference: HE and GREML (both implemented in GCTA, see Methods). Specifically, we examined how well the two methods inferred heritability across partitions of MAF when the true underlying heritability was known. We simulated a wide range of genetic architectures with heritability distributed across 8 MAF partitions using a sample size of 500 individuals and a total heritability of 0.8 (Simulation Set 3). We found that when n=500 individuals are simulated and analyzed using 8 MAF bins, GREML fails to converge ∼65% of the time, regardless of the fraction of heritability deriving from rare variants (Figure 2a).

**Figure 2.**
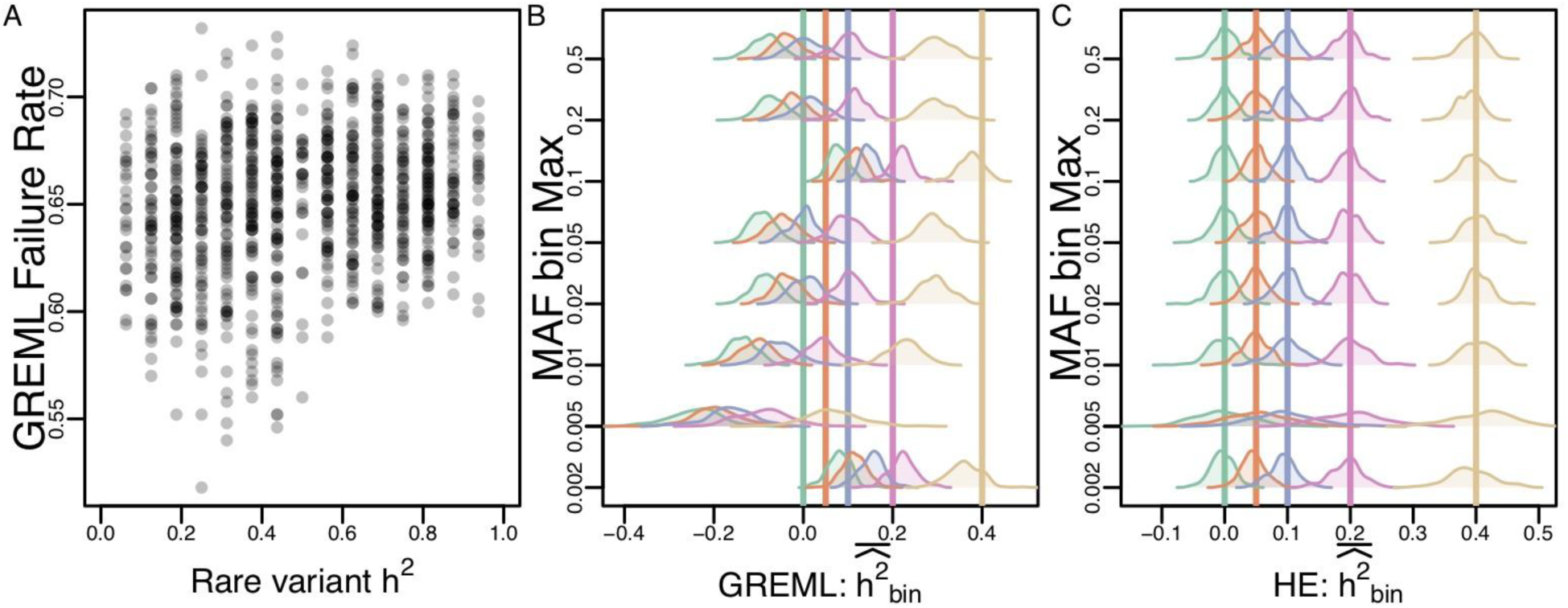
Simulations comparing GREML and HE. (A) The fraction of simulations that failed to converge as a function of the fraction of h^2^ that derives from rare variants (MAF<0.02). Each point represents 500 simulations of a different genetic architecture (see methods). For the GREML iterations that did converge, the distribution of mean h^2^ inferred across genetic architectures is shown for each MAF bin analyzed. True h^2^ shown as vertical bars. Similarly, (C) shows the distribution of mean h^2^ inferred for HE. Direct comparisons of point estimates and standard errors are shown in Figure S2.

When GREML does converge, the resulting heritability estimates can be biased (Figure 2b). In contrast, the regression framework of HE always provides a heritability estimate, and the inferred values tend to be unbiased under a broad range of conditions (Figure 2c). Figure S2 shows a direct comparison of heritability estimates across simulated parameters for the two algorithms and shows that the standard deviation of the heritability estimates across simulations tend to be comparable between HE and GREML.

Both HE and GREML report theoretical standard errors (SE) of the estimated heritability, but we found that neither algorithm report estimates of the SE that reliably reflected the empirical standard errors. While the SE reported by both algorithms are comparable for the higher MAF bins analyzed (MAF > 0.02), the reported SEs for the lowest MAF bin analyzed (0.001 ≤ MAF < 0.002) exhibit conflicting patterns (Figure S2).

When compared to the empirical standard error across simulations in a set, HE tends to grossly overestimate the SE of the estimate for the lowest MAF bin, while GREML tends to underestimate the SE of the estimate. As a result, approximate 95% confidence intervals 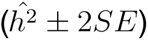 of the estimates for the lowest MAF bin are highly conservative for HE (100% of confidence intervals overlap the true *h*^2^) but become anti-conservative for GREML as the simulated *h*^2^ increases (only 83.8% of confidence intervals overlap the true *h*^2^ when the true *h*^2^ = 0.4; Figure S2). Given that HE tends to be less biased than GREML and not suffer from convergence issues, we exclusively used HE for the remainder of our analyses.

### Heritability Inference Quality as a Function of MAF Partitioning

Prior research has suggested that bias can be introduced when sites of differing MAF are pooled into the same GRM (Lee et al., 2013; Yang et al., 2015). We assessed this form of bias in a cohort of 500 individuals using heritability simulated across 8 MAF partitions (Simulation Set 3). We inferred the heritability of these simulated phenotypes either with the same 8 MAF partitions upon which they were simulated or pooled MAF bins (diagrammed in Figure 3a). The results of these inferences show that when variants are finely partitioned by MAF, the estimates are unbiased. As more of the MAF spectrum is included with the rarest partition, the estimate is upwardly biased by as much as 0.24 (30% of the total simulated heritability) when sites 0.001 ≤ MAF < 0.1 were pooled together. These biases in the total h^2^ estimates were driven by the estimates from the pooled variants, with the estimates from the remaining bins being relatively unbiased.

**Figure 3.**
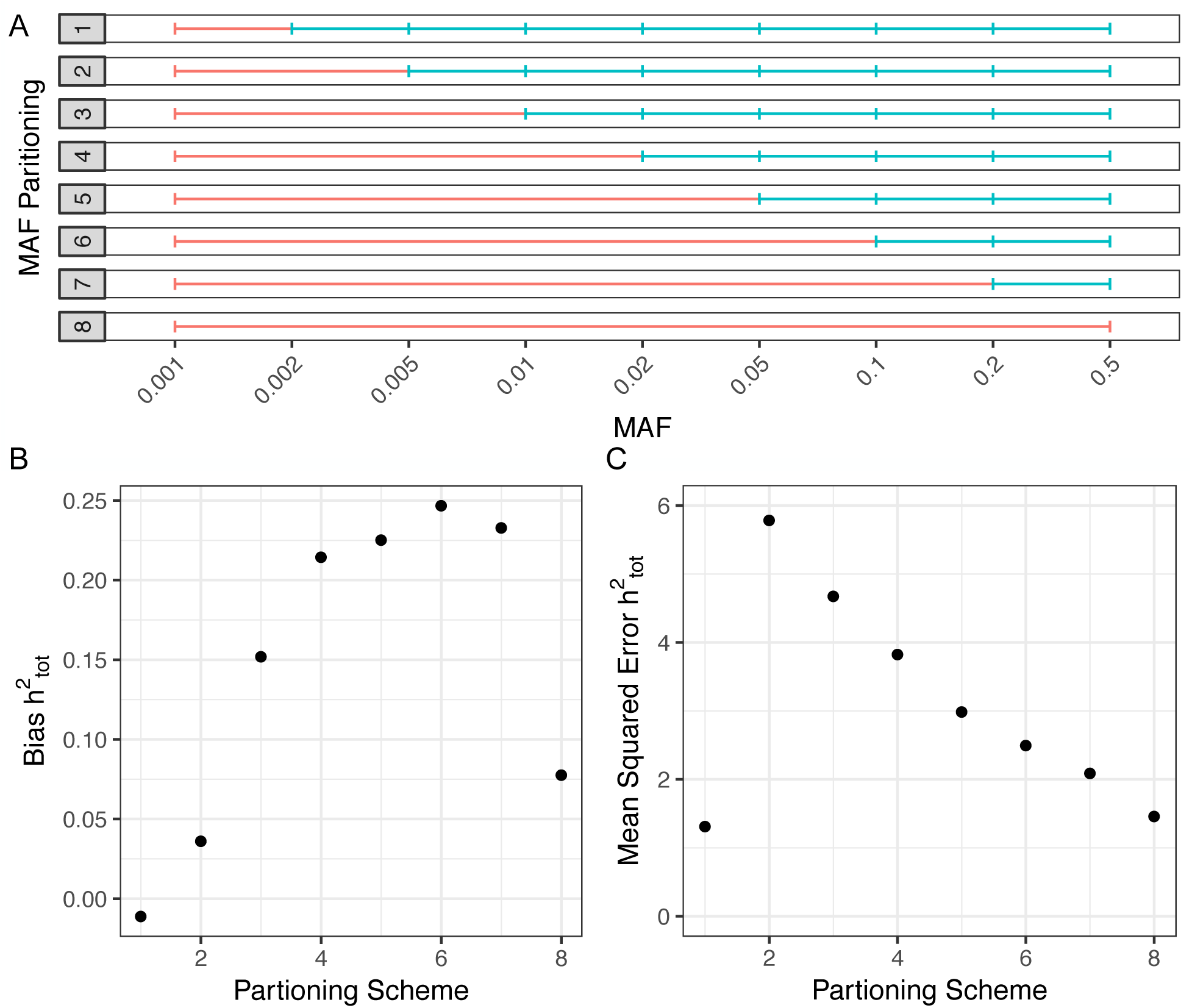
Impact of MAF Partitioning on Heritability Inference. (A) The partitioning scheme of the MAF spectrum. (B) Bias of the total inferred heritability for different partitioning schemes. (C) Mean squared error of different partitioning schemes.

Using the same set of simulations, we assessed the impact of pooling high and intermediate MAF partitions on the performance of HE regression (Figure S3 and Figure S4, respectively). We found that inference of heritability showed moderate downward bias when the highest MAF partitions are pooled, with the worst bias occurring when pooling MAF range (0.005, 0.5] with a bias of −0.08 (−10% of the total simulated heritability). Pooling variants of intermediate MAF resulted in less bias than the pooling of high MAF variants.

In any given study, issues of genotyping error, imputation, and MAF-dependent standard errors limit the lowest MAF that can be examined, and such sites are often excluded. We examined whether excluding the lowest MAF bins would bias the heritability estimates from the remaining MAF bins. We simulated phenotypes on 500 individuals using heritability distributed across 7 partitions (Simulation Set 2). We inferred heritability across the full 7 original partitions and successively excluding rare variants. The distributions of inferred heritability are shown in Figure 4. We found that exclusion of rare variants did not induce a bias in the estimates of heritability of the included bins, rather that the total estimated heritability would be an unbiased estimate of the variants that are included. As a result, any heritability attributed to the excluded MAF bins would simply remain as missing heritability.

**Figure 4.**
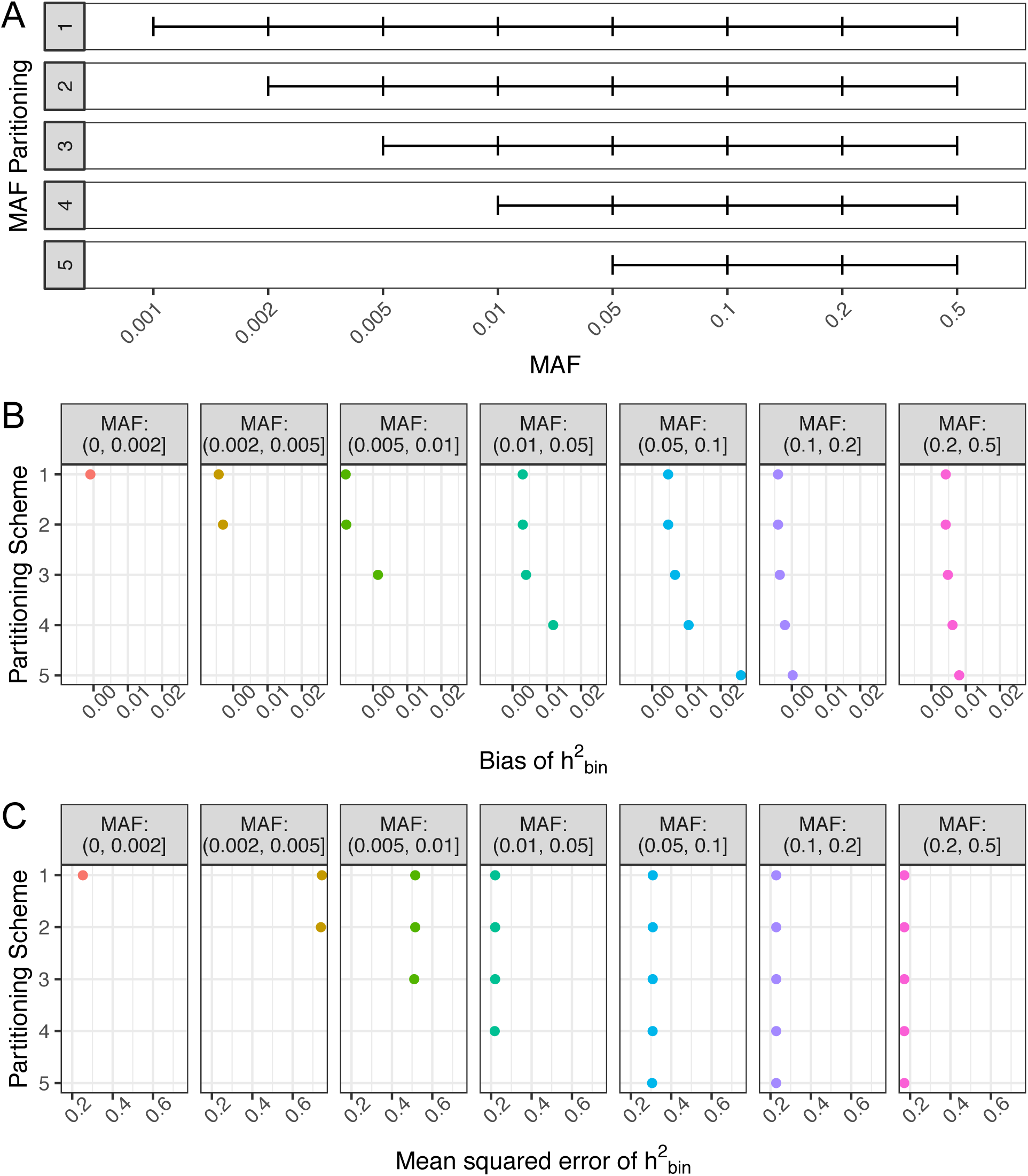
Impact of Excluding Low Frequency Variants on Heritability Inference. (A) The partitioning scheme of the MAF spectrum showing the exclusion of increasing range of the MAF spectrum. (B) The average bias of the inferred heritability of each partition included in the inference. (C) The mean squared error of the inferred heritability of each partition included in the inference.

### Impact of Sample Size on Heritability Inference Quality

The forces of natural selection will drive causal variants to different frequencies in the population. We sought to investigate how finely we can explore the population level MAF-heritability spectrum for different sample sizes. To this end, we simulated heritability partitioned across 34 LD-MAF partitions of quality-controlled, unrelated UK Biobank European population (17 MAF partitions each split by median LD score) on 50,000 individuals. We then inferred the heritability of these 34 partitions using the full cohort of 50,000 individuals, as well as subsets of 5,000 and 500 individuals. The magnitude of bias (Figure 5a) was generally larger for the lower MAF bins, and the scale of the bias was much higher for smaller sample sizes. Standard error (Figure 5b) generally increased for more rare partitions and decreased dramatically with larger sample sizes (dropping by more than a factor of 10 for each factor of 10 increase in sample size in many of the partitions).

**Figure 5:**
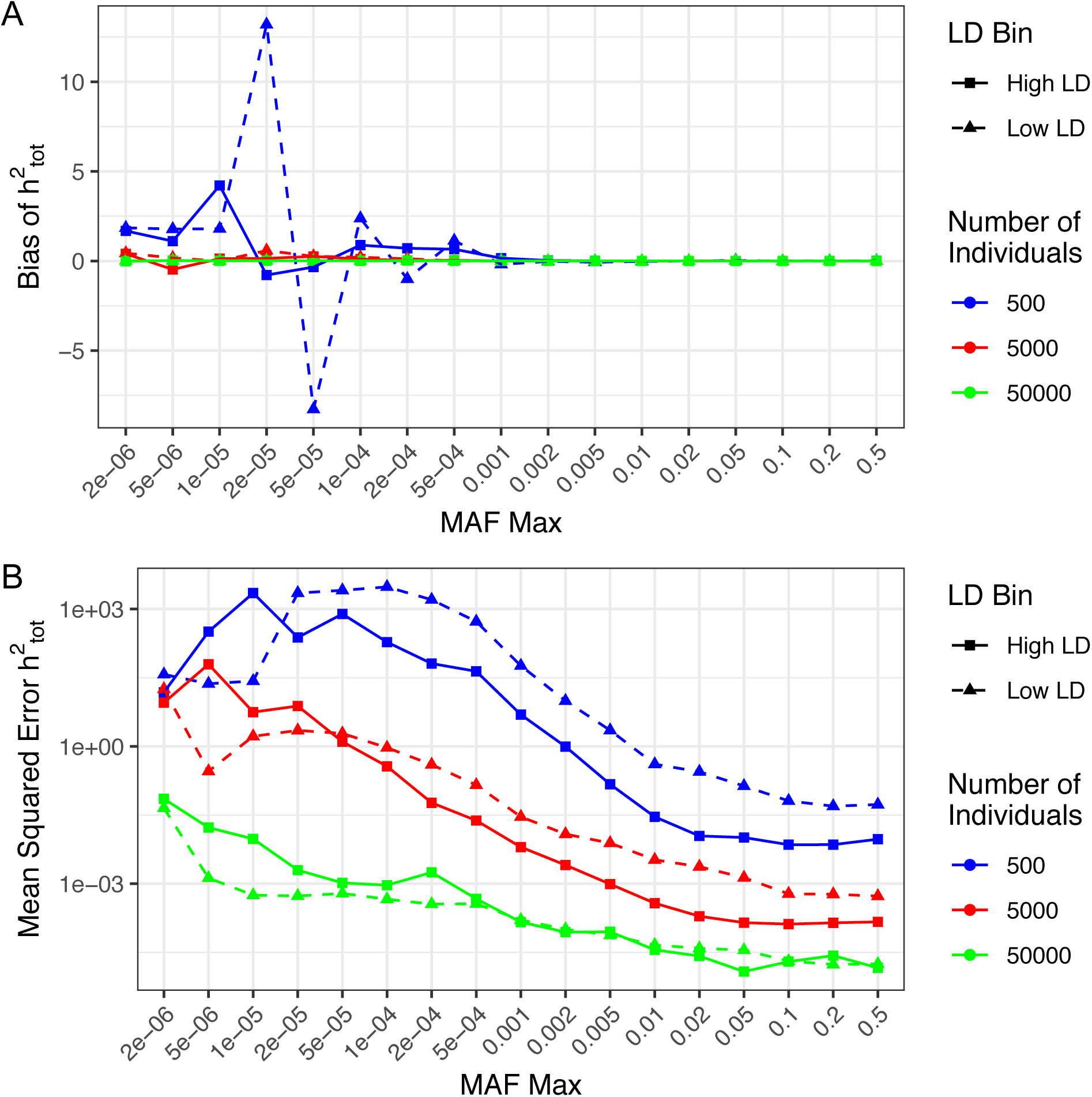
Impact of Sample Size on Bias and Mean Squared Error of Estimates. (A) Bias of inferred heritability with different sample sizes. (B) Mean squared error of inferred heritability with different sample sizes.

### Heritability of Complex Human Traits in the UK Biobank

We randomly selected 50,000 unrelated individuals to infer the genetic architecture of quantitative human traits. We restricted the analysis to the 72 quantitative traits among the biomedical categories blood, body, breath, and urine that were measured in at least 25,000 individuals. We used HE regression to infer the heritability of each trait using variants with MAF ≥ 1 × 10^−4^, partitioned across 11 MAF bins, each split into 2 LD bins (see Methods). To correct for population structure, we progressively added principal components (PCs) as covariates up to 15 PCs. We then added three geolocation covariates that describe where each individual lives (north/south, east/west, and distance from the coast; Figure S6). We found that there is only a subtle effect of adding additional PCs beyond the fifth PC. However, geolocation covariates corrected for an additional source of rare variant stratification (particularly for variants with low LD). For further analysis, we focus on the inclusion of 15 PCs and the three geolocation covariates.

The average total heritability of these traits was 0.269 (full list of 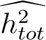 in Figure S5). Figure 6A shows the heritability estimates across MAF/LD bins. The plurality of heritability derives from the most common MAF bin (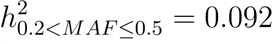, representing 34.3% of the average total heritability; Figure 6a). However, there is considerable variation in the contribution of different MAF bins to heritability of different traits (Figure 6B, which shows the cumulative, left, and reverse-cumulative, right, heritability across MAF bins for each of the 72 traits). Averaging across traits (Figure 6C), we find that little heritability derives from ultrarare variants. Superimposing the cumulative and reverse-cumulative heritability plots allows us to easily identify the MAF at which half the heritability has been described (the intersection of the cumulative and reverse-cumulative heritabilities). Overall, approximately half the heritability is explained by variants with MAF ≤ 0.05. Partitioning alleles by low versus high LD, we find that low LD variants constitute 3.3-fold more heritability than high LD variants, which is largely driven by low frequency variants (approximately half the heritability of low LD variants is explained by variants with MAF≤0.02), while heritability of high LD variants is primarily driven by common variants (approximately half the heritability of high LD variants is explained by the highest MAF bin alone).

**Figure 6:**
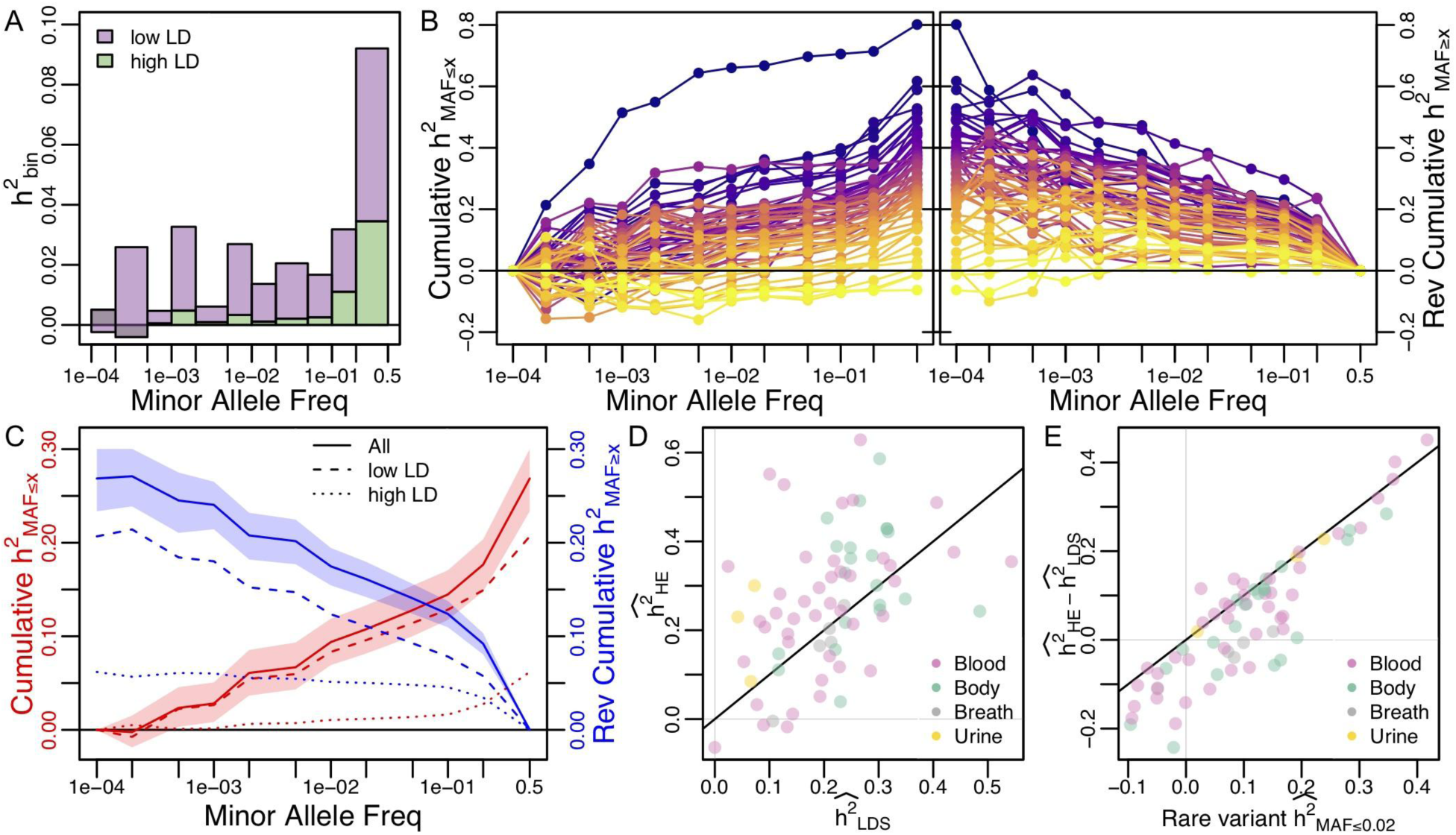
Heritability of Human Traits in UK Biobank. (A) Stacked bar plot of average heritability in each MAF-LD partition across 69 biomedical traits. (B) Cumulative and reverse cumulative heritability of all biomedical traits (with traits colored according to their total heritability, see Figure S5). (C) Average cumulative and reverse cumulative heritability across traits (solid line) with envelope showing the 95% quantile range from 1000 bootstrap samples. Dashed and dotted lines represent low and high LD partitions, respectively. (D) Comparison of the inferred total heritability across traits using HE regression (y-axis) versus LD Score (LDS) regression (x-axis). (E) Difference between HE and LDS heritability estimates versus our inferred rare variant (MAF≤0.02) heritability estimate. In D-E, points are colored according to the four biomedical categories of traits, with diagonal line show for reference.

Previous estimates of heritability from these data have been calculated using LD Score (LDS) regression (Walters et al., n.d.). Our estimates of total heritability using HE regression have a reasonable concordance with the LDS estimates (Figure 6D), with a correlation of *r*^2^ = 0.75. The discrepancies between our HE estimates and the LDS estimates are mostly driven by the contribution of low frequency variants (MAF ≤ 0.02) to our HE-based estimates (Figure 6e).

## Discussion

Our simulations show that drawing causal alleles and the effect sizes for those alleles independently of MAF will result in the majority of heritability arising from common alleles. While certain models could propose a relationship between probability of being drawn as a causal allele, effect size distribution, and minor allele frequency the actual relationship underlying actual traits remains unknown. If heritability inference procedures are tested and calibrated on a small subset of possible models, the performance on traits that do not fit that model may not be accurate. Indeed we found that REML exhibited substantial bias in many of our simulations. HE Regression, in contrast, was much more robust to a variety simulated heritabilities.

Our investigation into the performance of HE Regression underscored the importance of partitioning variant by MAF. The simulations we conducted also highlighted the importance of sample size in assessing the contribution of rare variants. A ten-fold increase in sample size reduced standard errors by more than a factor of ten for rare variants. The computational efficiency of HE Regression based methods should allow for examination of greater sample sizes, and therefore the examination of the contribution of rarer variants, as compared to REML.

Using a cohort of 50,000 individuals from the UK Biobank, we were able to examine the heritability of 72 biomedical traits down to a MAF of 0.01%. We found that these traits had average heritability was 0.269. Of this, 34.3% of the total heritability was found in the highest MAF partition and 34.9% of the total heritability was explained by variants with MAF ≤ 1%. These data are inconsistent with simulations that have independent and identically distributed effect sizes across MAF bins (where we inferred 67% of heritability to be due to the highest MAF bin; Figure 1). This suggests that causal variants are disportionately at low frequency or that these low frequency causal variants have larger effect sizes than common causal variants. The variants in regions of low LD accounted for 3.3-fold more heritability than those in regions of high LD, consistent with past findings (Zeng et al., 2018; Wainschtein et al., 2019) and is considered evidence of negative selection. That the variants with MAF≤0.02 explain roughly half of the heritability of the low LD variants may be further suggestive of negative selection acting upon the genetic architecture of these traits.

One important caveat to our analysis is that we have only considered variants identified through genotyping and imputing samples to an external reference panel. This means that a majority of ultrarare variants that are carried by the 50,000 individuals we studied were not included in our analysis. Indeed a recent study showed that there were more than ten times as many variants with MAF < 0.01% revealed through whole exome sequencing in a cohort of 50,000 UK Biobank individuals than in a genotyped and imputed comparable cohort (Hout et al., 2019). While rare-variant association studies are just as well powered with genotyped and imputed variants as they are with whole genome sequencing (Tong & Hernandez, n.d.), our ability to infer the contribution of these ultrarare variants to heritability of complex traits is nonexistent. While we did not conduct simulations directly to assess the impact of genotyping and imputation error, these effects would mostly be observed in the most rare MAF bins, where we only observed modest amounts of heritability. As technologies for collection of genetic material improve and computational feasibility of ever-larger cohorts is achieved, we will be better able to examine the contribution of ultrarare variants to heritability of human traits.

The findings here relate to the specific population studied, a non-random sample of the UK population. While findings may have some sensitivity to the inclusion of additional covariates, covariates must be examined on a case-by-case basis to avoid altering the interpretation of particular phenotypes. Future work can examine how these findings generalize to other populations.

## Acknowledgments

This research was undertaken, in part, thanks to funding from the Canada Research Chairs program (to R. D. H.) and by the National Institutes of Health (grant 1R01HG007644 to R. D. H.; grants R01CA201358 and R25CA112355 to J. S. W.) and by the Howard Hughes Medical Institute Gilliam Fellowship (to K. A. H.). We declare no conflicts of interest.

## Supplementary Materials

### Inclusion of Covariates

As GCTA has not implemented the inclusion of covariates in their HE Regression method, these were included as “pseudo GRMs.” Letting *c_i_* be the value of the *i*^th^ individual for the covariate C, the mean-centered, unit-variance-adjusted covariate, 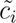, is:

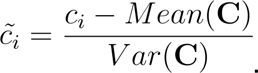

The entry of the covariate matrix for the pair of individuals *i* and *j*, Γ, would be 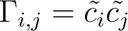. These covariate matrices were computed in Python and exported in a format matching that of GCTAs GRMs. Individuals missing values for covariates were replaced with median of the remaining values.

---

**Figure S1.**
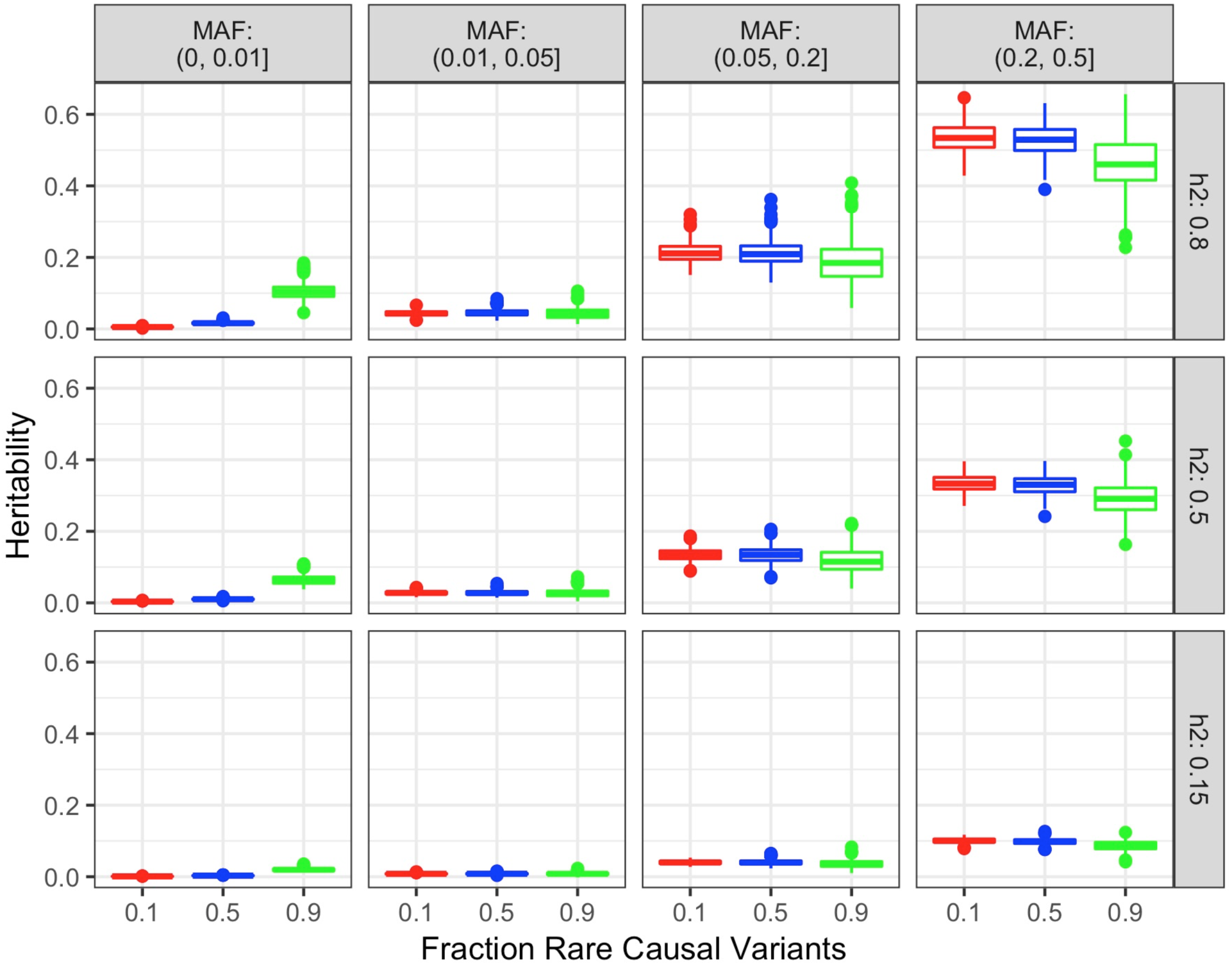
Distribution of Simulated Heritability Varying Fraction of Rare Causal Alleles across Different Total Heritabilities.

**Figure S2.**
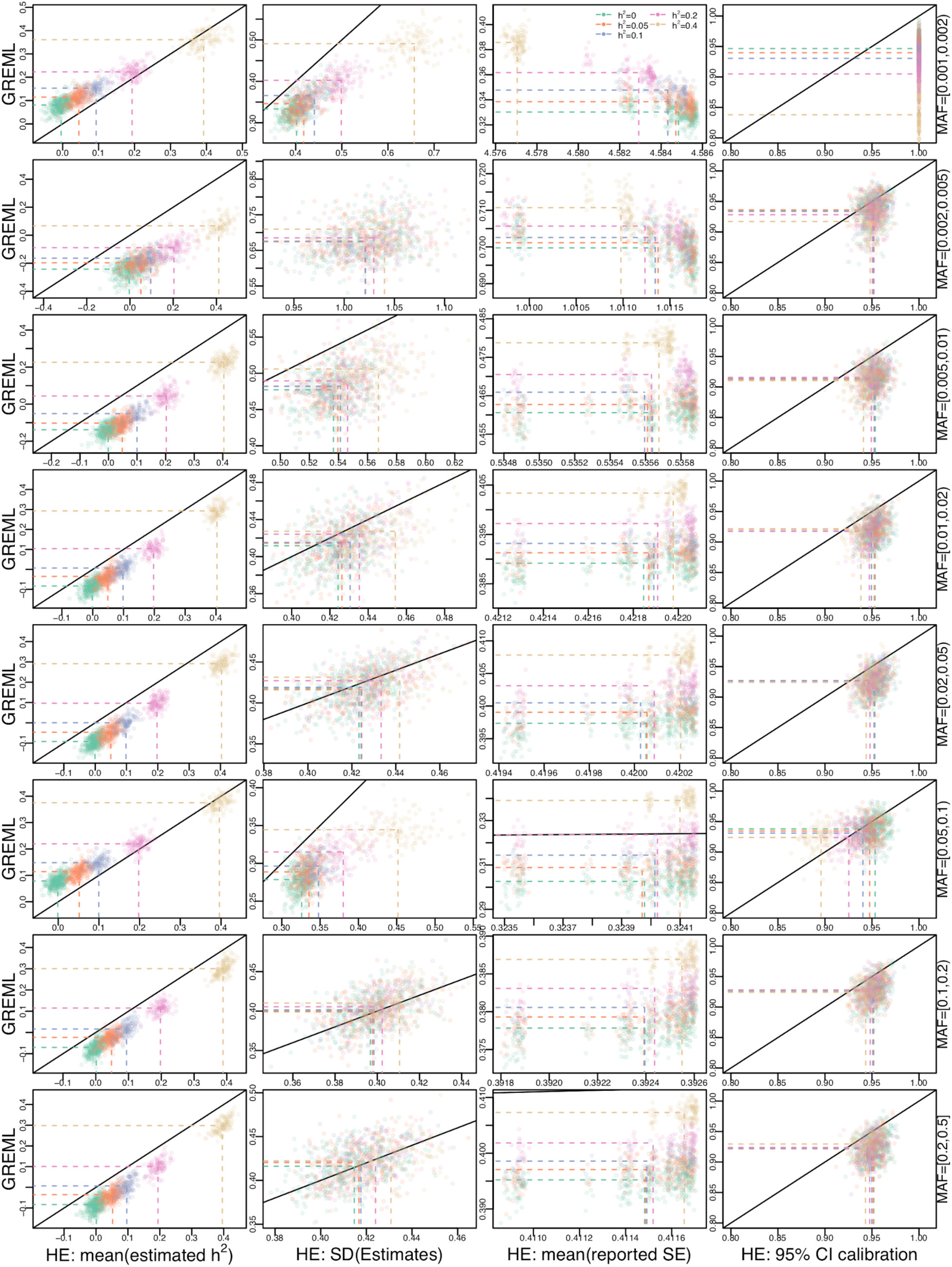
Simulations comparing GREML and HE. In all plots, each point represents 500 simulations of a single genetic architecture when the true total h^2^=0.8. Each row of figures represents a different MAF bin (rare variants at the top, common variants on the bottom), where each point is colored by the true h^2^ that derives from that MAF bin and is one of: green (h^2^=0), orange (h^2^=0.05), blue (h^2^=0.1), pink (h^2^=0.2), or brown (h^2^=0.4). Plots in the first column (left) compare the mean estimated h^2^ (across 500 simulations, or the number that converged, see main text Figure 2A) for GREML (y-axis) versus HE (x-axis). Note that the density functions in main text Figure 2B-C represent the marginal distributions of these points. The 2nd column of plots compare the standard deviation of the estimates for each genetic architecture. The third column of plots compare the reported standard errors from GREML vs HE. The fourth (right) column of plots compare the fraction of approximate 95% confidence intervals (CI) that overlap the true h^2^ for a given bin. In all plots, the dashed lines connect the average across all sets of simulations with the same true h^2^ in a bin to their axis, and the black line represents the y=x line.

**Figure S3:**
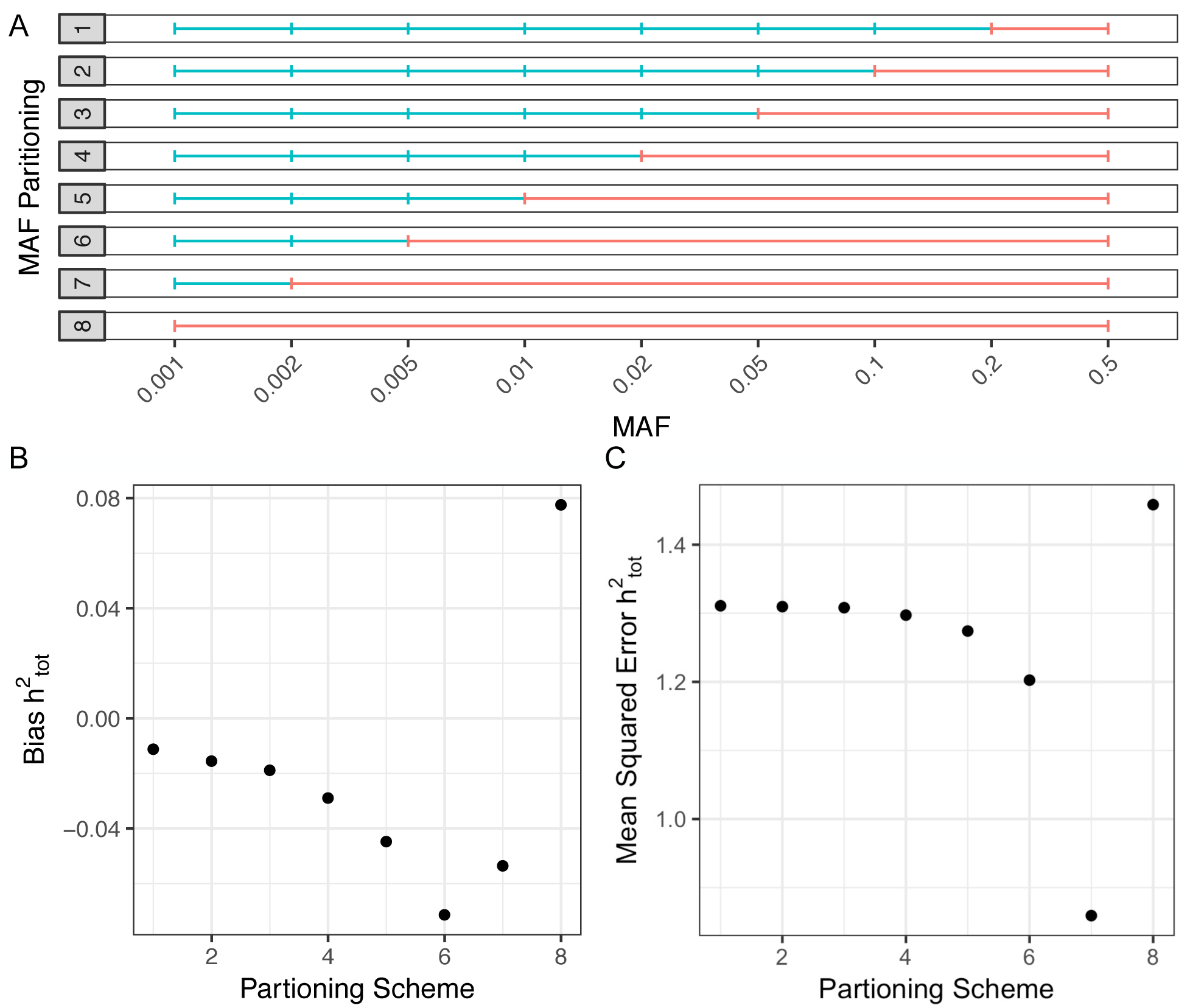
Impact of MAF Partitioning on Heritability Inference for High MAF. (A) The partitioning scheme of the MAF spectrum used for inference to investigate the impact of pooling variants of high MAF. (B) Bias of the total inferred heritability for different partitioning schemes. (C) Mean squared error of different partitioning schemes.

**Figure S4:**
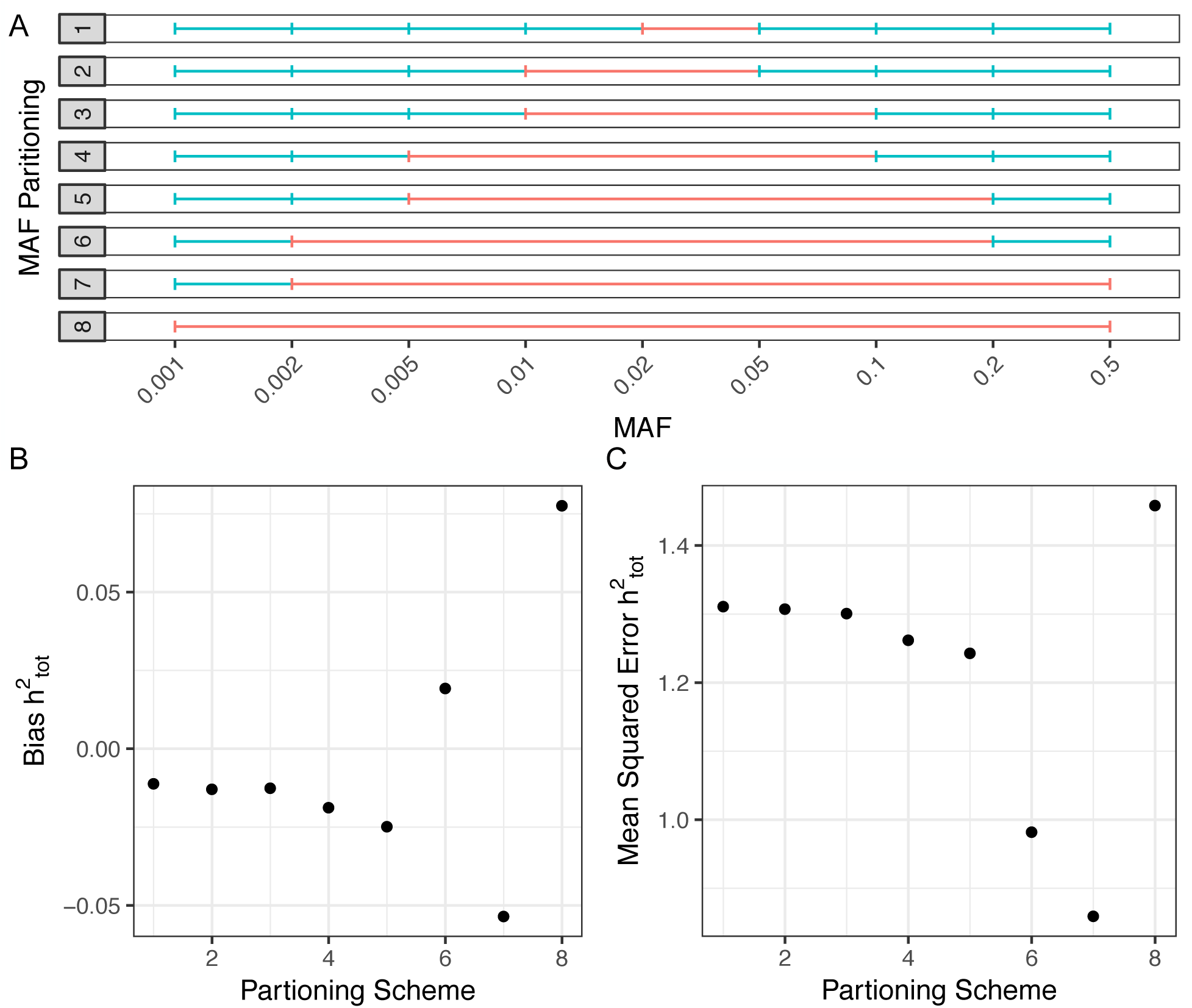
Impact of MAF Partitioning on Heritability Inference for Intermediate MAF

**Figure S5.**
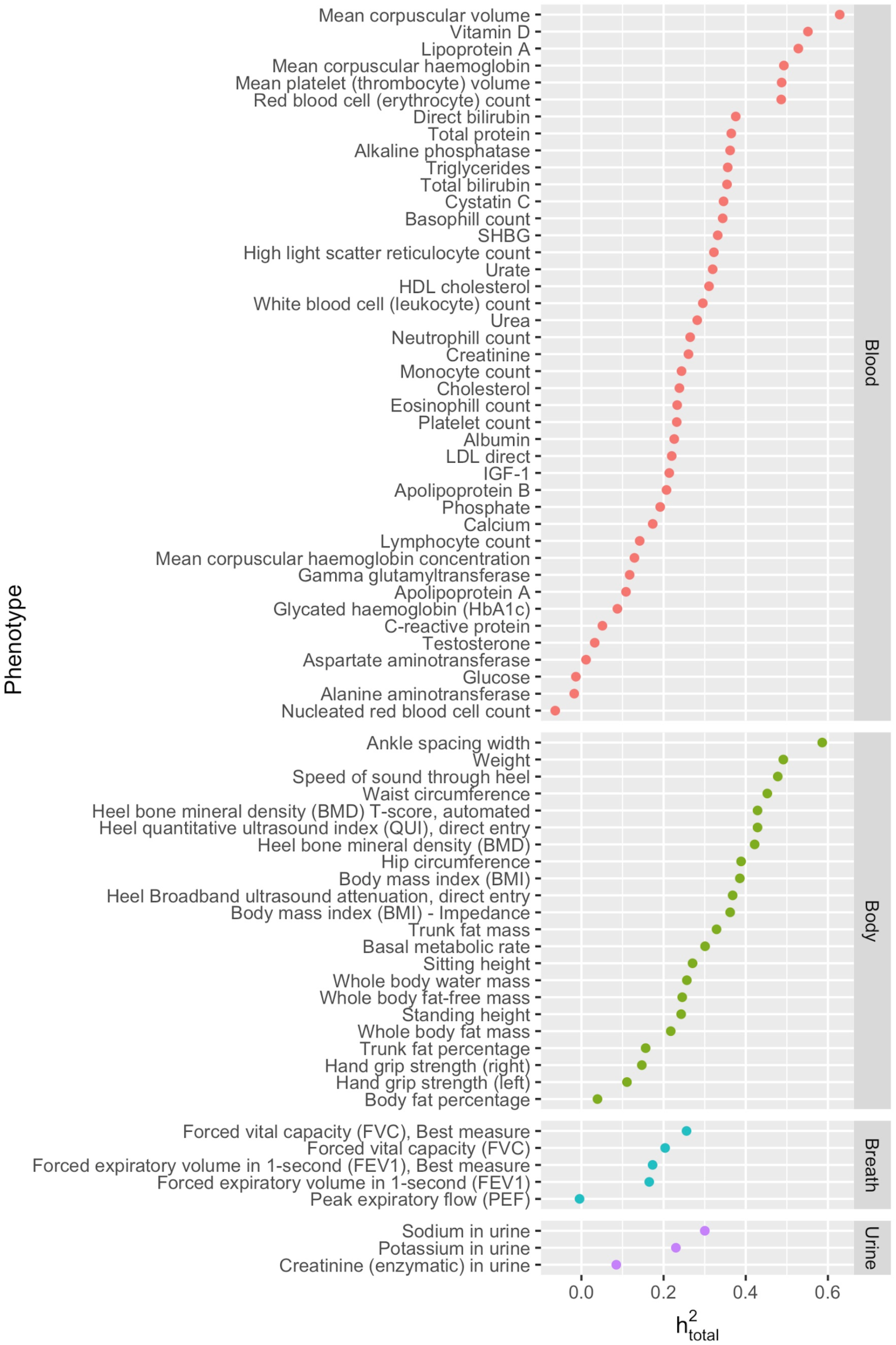
Inferred Total Heritability of Different Quantitative Measurements in UK Biobank. The total inferred heritabilities of the 72 biomedical traits examined.

**Figure S6.**
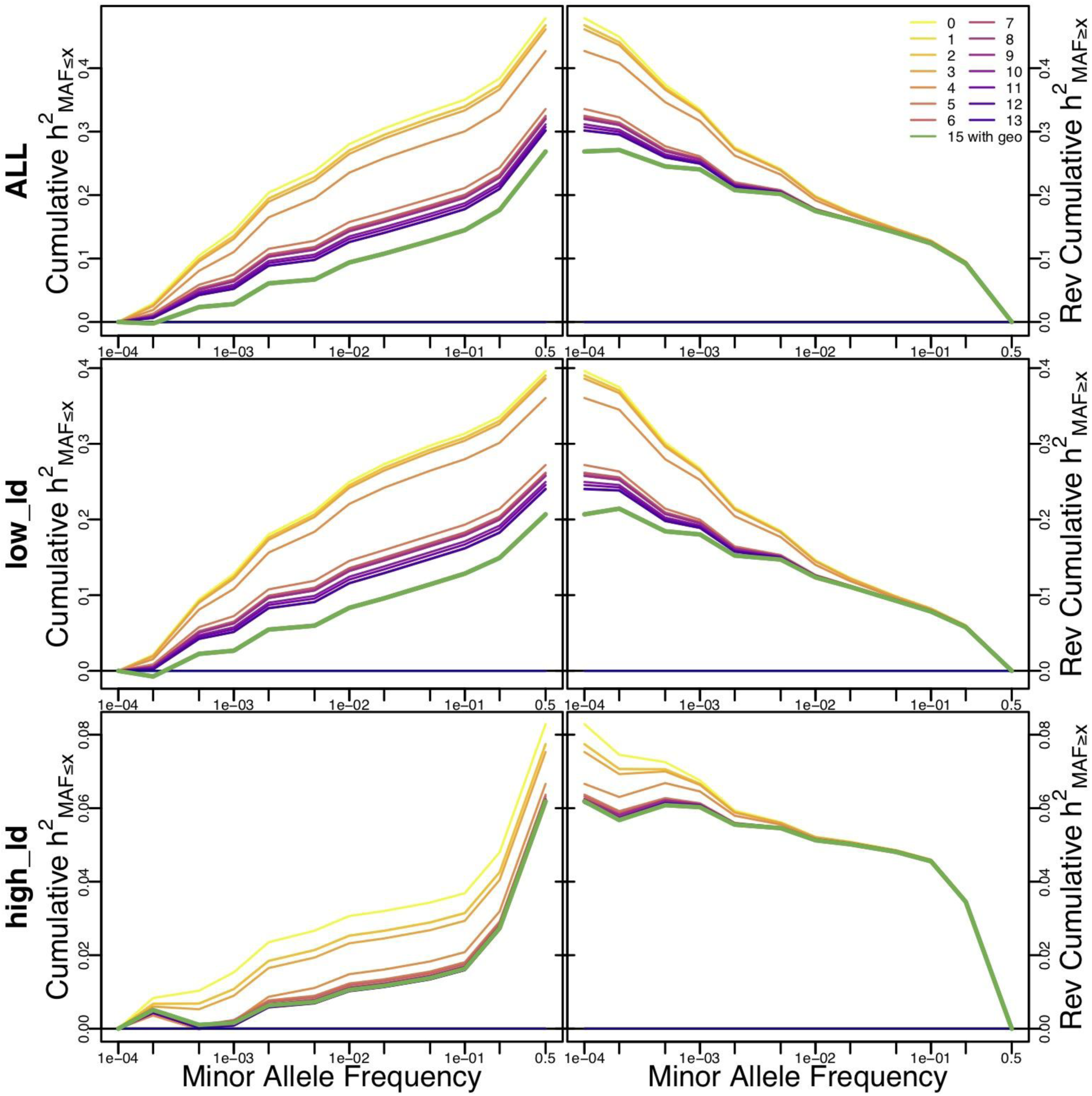
The genetic architecture of biomedical traits inferred with different covariates. The left panels show the cumulative heritability below a given MAF, and the right panels show the reverse cumulative heritability above a given MAF. Top panels show the average total heritability, while the middle and bottom panels examine the low LD and high LD bins (respectively).

